# Diversification of the “EDVID” packing motif underpins structural and functional variation in plant NLR coiled-coil domains

**DOI:** 10.1101/2025.06.01.657260

**Authors:** Oliver Sulkowski, Anna Ovodova, Andrea Leisse, Keelan Gögelein, Alexander Förderer

## Abstract

- Nucleotide-binding leucine-rich repeat receptors (NLRs) are critical in plant immunity and display remarkable allelic diversity. Coiled-coiled NLRs (CC-NLRs) are the most widespread group of these receptors found across flowering and non-flowering plants.
- Here we investigate the sequence conservation and functional variation of the conserved EDVID motif found in the α3-helix of the cell death inducing CC domain of plant NLRs. We analyse our findings in context of published protein structures and structure prediction.
- We find that the conserved EDVID motif can serve as a predictor of canonical CC-NLR function and oligomeric assembly.
- We also find that the EDVID motif is accompanied by preceding acidic residues in certain CC-NLRs with homology to the Arabidopsis CC-NLR RPP8. The appearance of this so-called preEDVID motif across the phylogeny of flowering plants and its contribution to the CC-NLR function underpins the structural diversity across NLRs with EDVID motif.
- We further show that CC-NLRs exist that have lost the EDVID motif sequence and function suggesting that this subgroup, previously referred to as CC_G10_-NLRs, functions in a different manner from the canonical mechanism.
- We find that acidic residues located to the α3-helix of the helper NLR NRG1.1 are linked to NRG1.1 cell death inducing activity.

## Introduction

Plants rely on a highly diversified innate immune system to defend against a broad range of pathogenic microbes. In turn pathogens secrete effector proteins into host cells to manipulate cellular processes and facilitate infection. In response, resistant plants recognise these effectors through a large family of intracellular immune receptors, the nucleotide-binding leucine-rich repeat (NLR) receptors inducing a programmed cell death at the infection site called hypersensitive response (HR) (Ngou *et al*., 2022; Jones *et al*., 2024). Incorporation of *NLR* genes into breeding programs has enabled durable, qualitative resistance against diverse crop pathogens. The ongoing evolutionary arms race between hosts and pathogens has driven extensive allelic diversity and gene family expansion in both plant NLRs (Martin *et al*., 2022) and pathogen effectors (Van de Weyer *et al*., 2019; Arroyo-Velez *et al*., 2020). However, most domesticated crops retain only a narrow subset of this natural NLR diversity.

Commonly plant NLRs feature a tripartite domain structure: an amino-terminal coiled-coil (CC), RPW8-like CC domain (CC_R_) or Toll/interleukin-1 receptor (TIR) domain, a central nucleotide and oligomerisation domain (NOD), and a carboxy-terminal leucine-rich repeat (LRR) domain. With the exception of non-singleton NLRs, this CC/TIR-NOD-LRR architecture combines recognition, regulatory and signalling function into a single amino acid chain. The NOD module allosterically links pathogen perception with immune signalling by the CC or TIR domain through regulatory adenosine nucleotide exchange. A recent burst in cryo-EM structures of plant NLRs has shown that they, much like their animal counterparts called inflammasomes and apoptosomes, assemble into oligomeric complexes upon activation, which are collectively referred to as resistosomes. While resistosomes formed by TIR-NLRs have enzymatic activity producing nucleotide-derived infochemicals and rely on the activation of the CC_R_-NLRs ADR1 and NRG1 (Huang *et al*., 2022; Jia *et al*., 2022), the CC-NLR resistosomes act as channels to release calcium from cellular storage as a second messenger (Förderer *et al*., 2022; Wang *et al*., 2019a; Bi *et al*., 2021; Jacob *et al*., 2021).

Phylogenetic classification of NLRs through their NOD module separates TIR and CC-NLRs (Chia *et al*., 2024) While TIR-NLRs with tripartite domain structure appear as an innovation of dicot plants (Johanndrees *et al*., 2023) CC-NLRs exist in monocots, dicots and in non-flowering plants (Chia *et al*., 2024). CC-NLRs contain several conserved motifs that enable motif based phylogenetic classification. Strongest conservation is present in the Walker A, also called P-loop, and Walker B motifs localised in the NOD module and mediating the binding and recognition of adenosine tri-and di-phosphates (Maruta *et al*., 2022). Specific amino acid substitutions in the Walker A and B motifs typically lead to a loss-of-function of the NLR due to a failure of nucleotide coordination in the binding pocket (Maruta *et al*., 2022). In contrast, single amino acid substitution in the conserved MH<u>D</u> (or sometimes VHD), motif for the sequence MH<u>V</u> (MHD^MHV^) lead to autoactivation of the NLR (Maruta *et al*., 2022). This autoactivation was shown to be linked to pathogen-independent autoimmunity phenotypes in plants (Tameling *et al*., 2006). Comparison of the protein structure of the ZAR1 NLR from *Arabidopsis thaliana* (Arabidopsis) in its inactive and active conformational states suggests that the MHD^MHV^ amino acid substitution stabilises ATP binding in the active, oligomeric form while destabilising the inactive NLR conformation (Wang *et al*., 2019a,b). The MHD^MHV^ autoactivating mutation, introduced to several NLRs has yielded cryo-EM structures and identification of NLR oligomeric state by blue native page of activated NLR complexes (Selvaraj *et al*., 2024; Liu *et al*., 2024; Tran *et al*., 2017)

Sequence conservation in the CC domain is overall weaker than in the NOD module (Sarris *et al*., 2016). Structure prediction using AlphaFold 3 has highlighted the conserved four-helix bundle architecture of the CC domain found in CC-NLRs and CC_R_-NLRs (Şulea *et al*., 2025). Upon activation of the NLR, the CC domain undergoes a structural rearrangement that leads to the protrusion of the α1-helix from the helix bundle. In the oligomer of ZAR1, this α1 helix forms a funnel-like structure that is thought to mediate insertion into the plasma membrane (Wang *et al*., 2019a; Bi *et al*., 2021). Insertion into lipid bilayer of the plasma membrane or other organelles necessitates the hydrophophic residues found in the α1-helix collectively referred to as MADA motif after the amino-terminal residues found in the CC-NLR *Sl*NRC4 of solanaceous plants (Adachi *et al*., 2019).

The most conserved and widespread sequence motif in the CC domain of plant NLRs is the EDVID motif (Glu-Asp-Ile-Val-Asp) (Wróblewski *et al*., 2018; Adachi *et al*., 2019). Previous investigation of this motif showed that it is present in 16% of the total 50 Arabidopsis CC-NLRs with the acidic aspartate and glutamate residues showing strongest conservation (Wróblewski *et al*., 2018) These acidic residues are also conserved in an additional 28% of Arabidopsis NLRs (Wróblewski *et al*., 2018). Structural investigation of the wheat NLR Sr35 has highlighted that the EDVID motif, which is presented as EDIVD in Sr35, forms four salt bridges to an arginine-cluster (R-cluster) in the LRR domain and that these R residues are spatially separated on the amino acid sequence by one iteration of the LRR-motif (Förderer *et al*., 2022) Sequence alignment with several NLRs from dicots and other monocots have shown that these arginines are co-occurring with the EDVID motif and is conserved in the functionally tested NLR Sr35 (Förderer *et al*., 2022). Although, the EDVID R-cluster ‘clamp’ is formed in both the inactive and active form of ZAR1, in the inactive AlphaFold 2 prediction of Sr35 and in the active cryo-EM oligomer structure of Sr35 (Wang *et al*., 2019a,b; Förderer *et al*., 2022) its function in the NLR activation mechanism is less understood. Mutations of the EDVID motif or R-cluster reduce or abolish cell death activity in Sr35 without negatively affecting NLR steady-state accumulation (Förderer *et al*., 2022)

Although several NLR structures have been elucidated in recent years (ZAR1 active and inactive, *Hv*MLA13, Sr35, *Sl*NRC4, *Nb*NRC4, *Sl*NRC2 dimer and filament, NRG1, NbRoq1, RPP1 etc.), they represent only a small subset of the thousands of plant NLRs. A comprehensive exploration of NLR structure and function is needed for agricultural applications that rely on the knowledge-base on the universality of these mechanisms and their variations across NLR subgroups, particularly the wide-spread CC-NLRs. Here we investigate the sequence conservation, structure and function of the critical EDVID motif mediating the single contact point of the LRR domain and the CC domain. Structural prediction of Arabidopsis CC-NLR oligomers shows two alternative EDVID motif positions in the protein structure: (1) a nested position in the oligomer with clearly defined R-cluster contact, or (2) a distal positioning of the EDVID motif with no R-cluster contact. To test the accuracy of these predictions, we made functional analysis in plants to test the different phylogenetic groups of CC-NLRs defined by Wróblewski et al. 2018. We used MHD to MHV amino acid substitution to probe for EDVID motif function. Different from effector-induced activation that is sensitive to EDVID mutation and results in reduced HR intensity, we found that in the autoactive substitution mutant D485V of Sr35, the cell death is not reduced by amino acid substitutions in the EDVID motif. Based on this finding we suggest that effector-induced activation may follow a different structural mechanism in this specific NLR. Using sequence alignments and this strategy of substitution in the MHD motif to trigger autoactivation, we found that the EDVID motif is present in 44% of Arabidopsis CC-NLRs and that its function is present in at least four NLRs from this phylogenetic subgroup. In the remaining 56% of Arabidopsis CC-NLRs and CC_R_ NLRs, strong EDVID motif conservation is not present, despite sporadic occurrence of acidic residues in α3-helix at the relative position of the EDVID motif. These CC-NLRs without a clear EDVID motif are unaffected in HR cell death activity upon amino acid substitutions. Interestingly, in the CC_R_ NLR NRG1.1 that is missing a clear EDVID motif, substitutions decrease HR cell death activity suggesting a critical role of these acidic residues in NRG1.1. Our results reveal functional sequence diversity in the EDVID motif of CC-and CC_R_-NLRs, emphasising that NLR structure and activation mechanisms differ between these subtypes.

## Materials and Methods

### Protein sequence alignments and structural prediction

Protein sequences were downloaded from the Arabidopsis Information Resource (TAIR) database (Arabidopsis.org, last use 27.04.2025)(Reiser *et al*., 2024) and aligned with MegAlign Pro (V. 17.5.0; DNASTAR Inc., WI, United States) using the default MUSCLE alignment algorithm. Structural prediction was performed using AlphaFold 3 (AF3) (alphafoldserver.com, last use 27.04.2025) (Google LLC, California, United States). To generate the active conformational state AlphaFold was set to generate the NLR oligomer (5 proteins) together with the ligands ATP (5x) and 50x palmitic acid to simulate a membrane (Madhuprakash *et al*., 2024). Structural alignment was performed with PyMOL (Schrödinger LLC, New York, United States) and molecular graphics where generated with UCSF ChimeraX 1.8 (Pettersen *et al*., 2021). AF3 structures were uploaded to following cloud server: https://nextcloud.mpimp-golm.mpg.de/s/sdydCNQTbYBJ9b7

### Hypersensitive response assays and Western blot analysis

For the transient gene expression, the synthesised constructs were cloned into the pDONR207 vector (Invitrogen) and mutated through Gibson assembly. To bypass the need to activate each NLR with their specific pathogen effector, where most of them are still unknown, mutations of the MHD-motif were generated to turn NLR autoactive. The generated plasmids were then recombined by an LR clonase II (Thermo Fisher Scientific) reaction into *pGWB517-4xMyc* with a C-terminally fused 4xMyc epitope tag(Nakagawa *et al*., 2007a). AvrSr35 was recombined into *pGWB411-FLAG* with a C-terminally fused FLAG epitope tag(Nakagawa *et al*., 2007b). The constructs were verified by Sanger sequencing and further transformed into *Agrobacterium tumefaciens* GV2260 by electroporation. The transformants were grown on YEB media selection plates containing rifampicin (50 mg mL^-1^), carbenicillin (100 mg mL^-1^) and spectinomycin (75 mg mL^-1^). A streak of transformants was picked and cultured in YEB medium with the corresponding antibiotics at 28 °C for 16 h, then harvested at 1681-times g for 10 min. The pellet was resuspended in infiltration buffer containing 10 mM MgCl2, 10 mM MES pH 5.6 and 100 µM acetosyringone and incubated in darkness at 28°C for 1h at 200 rpm. For Western blot analysis all OD_600_ was adjusted to 1.0. For the HR-assays the NLR and their respective amino acid substitution mutants had the following OD_600_: Sr35 (0.06), RPM1 (0.3), RPP8 (0.6), RPP13 (0.6), *Bj*WRR1 (0.2), SUMM2 (0.2), L (0.05) and NRG1.1 (0.3). For the effector-based activation, Sr35 was co-infiltrated with AvrSr35 with OD_600_ 1.0 and in the negative control with empty vector OD_600_ 1.0. Infiltration was performed with a sterile 1 mL syringe to mature leaves of five week old *Nicotiana benthamiana*.

Phenotypic data was documented 4 days after infiltration through photographs and infrared scans with an excitation range of 745 to 765 nm, an emission range of 800 to 850 nm, and three seconds exposure time with the iBright® gel documenting system (Thermo Fisher Scientific Inc, Darmstadt, Germany) (Xi *et al*., 2021). The resulting black and white scans were analysed with the image processing software Fiji (v. 2.16.0) (Schindelin *et al*., 2012) through measuring the mean grey value (M.G.V.) of the infiltrated area. For the Western blot analysis, the infiltrated leaf discs were harvested 24 h after infiltration, flash-frozen in liquid nitrogen and ground to powder using a Retsch grinder. The plant powder was mixed with Urea/SDS-buffer (8M Urea, 2% SDS, 20% Glycerol, 100 mM Tris-HCL pH 6.8, 10 mg BPB, 100 mM DTT) in a 1:2 ratio and boiled at 95 °C for 5 min. 3 µL were loaded onto 10% SDS-PAGE. The protein was transferred over night at 4 °C to a PVDF membrane, which was blocked for 2 h with 5% fat-free milk powder and detected using monoclonal mouse anti-MYC antibodies (1:3,000; Novex) with overnight incubation at 4 °C and polyclonal goat anti-mouse IgG-HRP (1:7,500; Abcam AB6728, Charge:1046144-19) antibodies with incubation at room temperature for 3 h. Protein was detected using SuperSignal West Femto substrates (Thermo Fisher Scientific Inc, Darmstadt, Germany) in a 1:1 ratio.

### Statistics

The HR-assay data was analysed in R-Studio (R v. 4.4.1) and displayed as box plots with the median and interquartile range. Normality was tested using the Shapiro-Wilk test and significance was tested with ANOVA and Tuckey HSD post-hoc test for pairwise comparison. Different letters represent significantly different groups of means, based on analysis of variance ANOVA and post-hoc Tuckey’s test P=<0.05.

## Results

The EDVID motif in the α3-helix of the CC domain of NLRs is a conserved sequence mediating the only intramolecular contact packing the ligand-binding LRR domain to the signalling CC domain in Sr35 (figure 1, a) and ZAR1. To investigate its role in the protein structure context of other NLRs, we performed structure prediction of RPM1, RPP8, SUMM2 and NRG1.1 oligomers using AlphaFold 3 multimer (figure 1, b). We chose these CC-NLRs based on a previous study by Wróblewski et al. 2018 that classified all 50 Arabidopsis CC-NLRs according to their NOD module and CC domain and that defined five distinct phylogenetic NLR groups varying in their EDVID motif: group A that represents the helper CC_R_ NLRs NRG1.1 without a clear EDVID motif (figure 1, b), SUMM2 from group B that contains additional NLRs without clear EDVID motif conservation also called CC_G10_-type NLRs (Madhuprakash *et al*., 2024) (figure 1, b), RPM1 from group C representing the canonical EDVID motif group with NLRs of the ZAR1-type and clear motif conservation (figure 1, b), and RPP8 from the phylogenetically separated group D that also shows clear EDVID conservation with additional acidic amino acids preceding it (figure 1, b). Group E of Wróblewski et al. 2018 containing eight of the Arabidopsis NLRs were excluded from our analysis due to the lack of CC domains or NOD modules in NLRs of this group. Predicted structures of RPM1 and RPP8 both with a conserved EDVID showed a distinctly nested placement of the CC domain barrel inside the complex with a clear prediction of salt bridges formed between acidic residues of the EDVID motif and basic residues such as arginines of the LRR domain (figure 1, b). This striking similarity of EDVID placement in RPM1 and RPP8 with Sr35 and ZAR1 is also accompanied by high ipTM and pTM confidence values, pLDDT scores above 70, of the AlphaFold 3 prediction and a small expected position error for the residues forming the intramolecular contact (figure 1, b, c). In contrast, oligomer prediction of NRG1.1 and SUMM2 that have no clearly conserved EDVID showed a strikingly protruded placement of the CC domain barrel outside of the complex (figure 1, b). Notably, this CC barrel protrusion is formed from the opposite end of the oligomer containing the NBD domain (figure 1, b). This alternative topology of the CC domain predicted by AlphaFold 3 was also accompanied by a lack of intramolecular contacts mediated by the α3-helix of the NRG1.1 and SUMM2 CC domains and the lateral side of the LRR domain carrying basic amino acids in Sr35 and ZAR1 (figure 1, b). Solvent exposure of these residues also yielded a low ipTM, pTM and pLDDT below 70 in the structure prediction (figure 1, b inlets) and low scores in the expected position error matrix (figure 1, c). These data suggest that while the CC domain of NLRs lacking conservation in the EDVID motif, such as SUMM2 and NRG1.1, may function in a different manner, the CC domain of NLRs with a conserved EDVID motif may function in the same way as ZAR1 and Sr35 in the oligomer.

**Figure 1.**
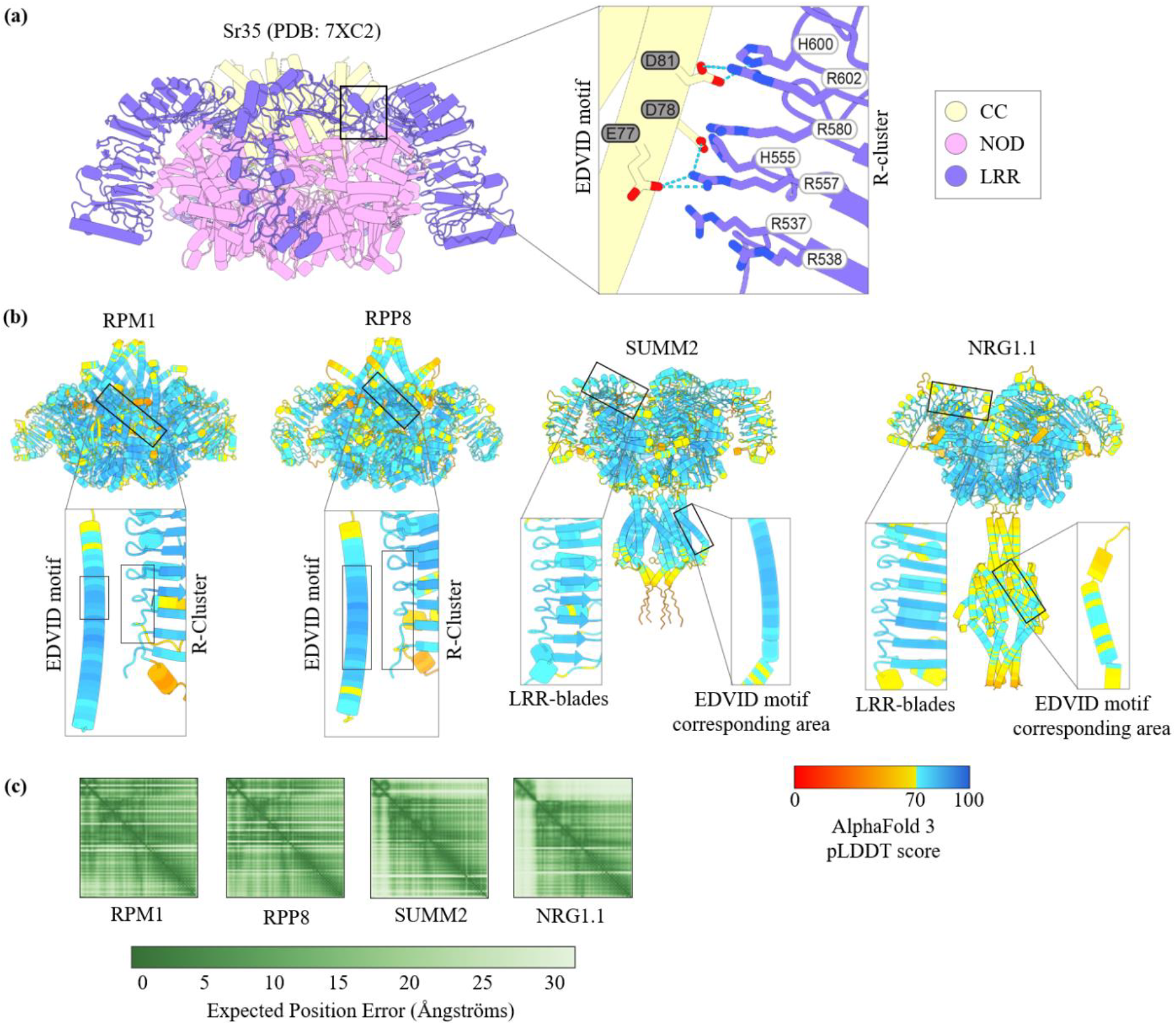
Oligomer of Sr35 and AlphaFold 3 predictions. **(a)** The cryo-EM structure of the Sr35 pentamer (PDB: 7XC2) displays saltbridges between the acidic amino acids of the EDVID (Glu, Asp, Val, Iso, Asp) motif on the coiled coil (CC) domain and basic amino acids of the arginine (R)-cluster on different blades of the leucin rich repeat (LRR) domain bridging the nucleotide binding and oligomerisation domain (NOD) module. **(b)** The AlphaFold 3 predictions of the Arabidopsis NLR RPM1 (ipTM=0.77; pTM=0.79) and RPP8 (ipTM=0.71; pTM=0.73) also display saltbridges while the predictions for SUMM2 (ipTM=0.60; pTM=0.63) and NRG1.1 (ipTM=0.57; pTM=0.60) place the domains apart. **(c)** The expected position error matrices of the first protomer of the predicted structures suggest alterative placements of SUMM2 and NRG1.1 the CC domains.

To test EDVID motif function across the four Arabidopsis CC-NLR groups and to test the merit of the AlphaFold 3 structure predictions, we did functional experiments *in planta*. Due to the diversity of potential activation triggers such as direct or indirect activation mechanisms, paired NLR mechanisms and others, we decided to activate NLRs of different groups in the same manner by introducing the autoactive MHD^MHV^ amino acid substitution. Starting with Sr35 of wheat we measured the effect of MHD^MHV^ - versus effector-mediated activation (figure 2, a). As expected, when wildtype (WT) Sr35 was transiently expressed in *Nicotiana benthamiana* (Nicotiana) by Agrobacterium-mediated transformation, only a slight HR phenotype could be observed through the overexpression of the cell death conferring receptor comparing to the empty vector (EV) (figure 2, a). Upon co-expression with AvrSr35, the WT Sr35 receptor produced a significantly stronger cell death (figure 2, a). Reproducing our previous finding (Förderer *et al*., 2022), the activity of Sr35 was still significantly less despite a reduction in the amino acid substitutions in Sr35 at the EDVID motif, from Y74A E77A D78A D81A in our previous study to E77A D78A D81A (figure 2, a). On the other hand, when the same EDVID motif amino acid substitutions E77A D78A D81A were introduced in a version of Sr35 that was autoactivated through the amino acid substitution MHV (henceforth called MHD^MHV^ ΔEDVID double motif mutant), this EDVID motif mutant displayed an HR phenotype similar to the Sr35 MHD^MHV^ autoactive mutant (figure 2, a). Western blot analysis confirmed the expression of NLR constructs in Nicotiana. Notably, the autoactive MHD^MHV^ mutant was not detectable, whereas the corresponding EDVID mutants in the same background showed clear expression. This likely reflects reduced steady-state protein levels of the autoactive NLR due to membrane association of the oligomerised complex and/or cell death associated degradation or suppression of expression (figure 2, b). These data show that effector-mediated activation of Sr35 is perturbed by EDVID mutation, whereas MHV-mediated activation is insensitive to mutation of the EDVID motif. This suggests that activation of Sr35 by MHD mutation is sufficient and independent of intramolecular stabilisation of the CC domain by the LRR domain. On the other hand, activation of Sr35 through binding of AvrSr35 to the Sr35 LRR domain may require the fixed topology of the LRR domain mediated by the EDVID motif.

**Figure 2.**
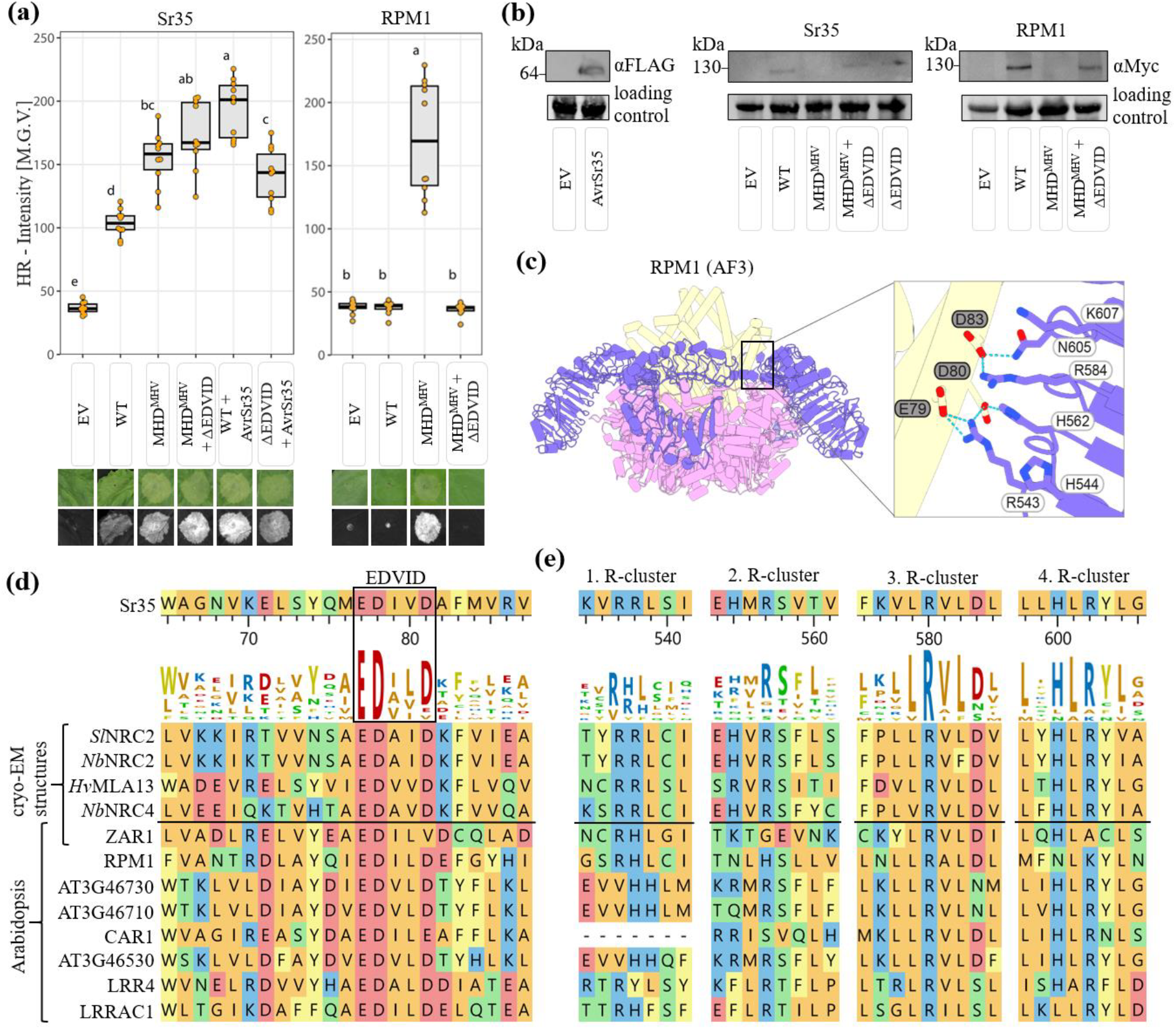
The EDVID motif is present across phylogenetically divergent species and can serve to predict canonical resistosome function. **(a)** Hypersensitive response intensity of 5 weeks old *Nicotiana benthamiana* measured through infrared fluorescence at 800 nm with the iBright® imaging system. Displayed are the mean grey values of the cell death measured 4 days post infiltration with *Agrobacterium tumefaciens*. Substituted amino acids for Sr35 and RPM1 are listed in table 1. The box plots display the median and interquartile range; n=10; different letters represent significantly different groups of means, based on analysis of variance ANOVA and post-hoc Tuckey’s test P=<0.05. **(b)** Western blot of α-Myc tagged NLR constructs expressed in *N. benthamiana*. Three replicates were pooled. Rubisco band on the blotting membrane as loading control. **(b)** Salt bridges between the acidic residues of the EDVID motif and the R-cluster on the LRR-domain in RPM1 (AF3 prediction ipTM=0.77; pTM=0.79). **(d)** Alignments of cryo-EM solved CC-NLRs (PDBs of *Sl*NRC2: 8XUV; *Nb*NRC2: 8RFH; *Hv*MLA13: 9FYC, *Nb*NRC4: 9CC8) with Arabidopsis CC-NLR show a conserved EDVID motif.

To study the functional relationship of the EDVID motif and the MHD motif and whether the fixed LRR-CC topology of Sr35 is required for activation of canonical EDVID NLRs, we introduced corresponding motif mutations in the Arabidopsis NLR RPM1 (figure 2, c). Comparing the RPM1 MHD^MHV^ motif mutant with an RPM1 MHD^MHV^ ΔEDVID double motif mutant, we observed that the expression of the double motif mutants of this receptor displayed a reduced HR phenotype compared to the single MHD motif mutant (figure 2, a). This suggests that the active form of RPM1 stabilised by the MHD mutation critically depends on a fixed CC-LRR domain topology for mediating cell death. This contrasting behaviour of RPM1 to Sr35 may be reflective of its different activation mechanism that depends on binding of the phosphorylated host protein RIN4 or the localisation of RPM1 to the plasma membrane in the inactive form (Chung *et al*., 2011). In Sr35, additional stabilizing interactions, such as intraprotomer or NOD-CC contacts, may help to lock the CC domain within the oligomer, whereas in RPM1, the EDVID-LRR interaction may play a more central role in enabling membrane insertion and cell death. Thus, despite sharing a conserved EDVID motif, group C NLRs may realise similar oligomeric topologies through different structural reinforcements. Our findings highlight previously unappreciated structural and mechanistic diversity within this NLR subclass. To support these functional differences with structural evidence, we aligned canonical EDVID Arabidopsis NLRs with structurally resolved NLRs from other species (SlNRC2, NbNRC2, HvMLA13, NbNRC4) (Ma *et al*., 2024; Madhuprakash *et al*., 2024; Lawson *et al*., 2025; Liu *et al*., 2024). The EDVID motif and adjacent R-cluster are highly conserved, underscoring their structural relevance across species (figure 2, d, e).

**Table 1.** List of all amino acid substitutions in the NLRs used in this study.

| NLR | MHD<sup>MHV</sup> | MHD<sup>MHV</sup> +<br>ΔEDVID |  |  |
| --- | --- | --- | --- | --- |
| NRG1.1 | D485V |  | MHD<sup>MHV</sup> + ΔEDVID 1:<br>D485V D63A E68/E71A | MHD<sup>MHV</sup> + ΔEDVID 2:<br>D485V D74/80A E77A |
| SUMM2 | D478V | D478V<br>E83/88A |  |  |
| L5 | D480V | D480V<br>D87/89A |  |  |
| Sr35 | D503V | D503V E77A<br>D78/81A | WT + ΔEDVID:<br>E77A D78/81A |  |
| RPM1 | D505V | D505V E79A<br>D80/83A |  |  |
| RPP8 | D494V | D494V<br>E73/77A D74A | MHD<sup>MHV</sup> + ΔpreEDVID:<br>D494V E63A D64/67A |  |
| RPP13<br>(AT1G59218) | D500V | D500V<br>E73/77A D74 | MHD<sup>MHV</sup> + ΔpreEDVID:<br>D500V E63A D64/67A |  |
| BjWRR1 | D490V | D490V<br>E73/77A D74A | MHD<sup>MHV</sup> + ΔpreEDVID:<br>D490V E63A D64/67A |  |

We next examined a subset of Arabidopsis NLRs that exhibit additional acidic residues N-terminal to the conserved EDVID motif. Sequence alignment revealed a conserved amino acid pattern preceding the EDVID motif, which we refer to as the preEDVID (figure 3, a). Similar to the ‘– – F F – ‘pattern of the canonical EDVID motif, this preEDVID motif follows a ‘– – Φ + – ‘property pattern, where ‘–’ indicates acidic residues (aspartate or glutamate), ‘+’ stands for basic residues (lysine, arginine or histidine) and ‘Φ’ denotes hydrophobic residues. Arabidopsis RPP8 homologs overall are highly conserved in the amino acids surrounding the EDVID and preEDVID compared to other Arabidopsis NLR groups (figure 3, a). To study if the preEDVID has stronger conservation than the surrounding amino acids, we included NLRs from a blast result using the preEDVID plus EDVID motif region from other Brassicaceae (*Brassica juncea, Capsella rubella*,), Malvaceae (*Hibiscus syriacus*), Vitaceae (*Vitis vinifera*), Lauraceae (*Persea americana*), monocots (*Oryza sativa*), and ANA grade species (*Nymphaea colorata, Magnolia sinica*) in the sequence alignment. Although *Bj*WRR1 has high similarity to the Arabidopsis NLRs, the alignment with other species showed that the preEDVID motif and the corresponding R-cluster has higher conservation than the diversified amino acids in the surrounding (figure 3, a, b) across a wide selection of flowering plants indicating a disperse appearance of the preEDVID motif in flowering plants. To study the function of the preEDVID motif in a protein structure context, we mapped its location in the RPP8 oligomer predicted by AlphaFold 3. This analysis revealed that both the EDVID and the preEDVID motifs are located on α3-helix of RPP8 and that the acidic residues of the motif form several salt bridges with the corresponding basic residues on the LRR domain (figure 3, c). To test if the EDVID and preEDVID act together in stabilizing the CC-LRR contact in RPP8 and RPP13, we did functional experiments in Nicotiana. Comparing the preEDVID motif mutation with the EDVID motif mutation in a MHV autoactive NLR background of RPP8 and RPP13, we found that EDVID mutation of these NLRs resulted in a complete suppression of cell death. On the other hand, mutation of the preEDVID resulted in an intermediate reduction of cell death compared to the MHD^MHV^ and the MHD^MHV^ ΔEDVID double motif mutant (figure 3, d). To study if the non-Arabidopsis NLR *Bj*WRR1 showed similar activity, we included it in the functional analysis (figure 3, e). The WT of *Bj*WRR1 showed induction of cell death and this activity was not increased by introduction of the MHV amino acid substitution (figure 3, e) suggesting that a missing negative regulator in the Nicotiana genetic background that controls *Bj*WRR1 induced cell death in *Brassica juncea*. To test the function of the EDVID and preEDVID motifs, we introduced the motif mutations in the *Bj*WRR1 MHV motif mutant. As in RPP8 and RPP13, EDVID motif mutation resulted in a complete loss of cell death induction, whereas the preEDVID mutation in *Bj*WRR1 did not result in a significant reduction (figure 3, d, e). Western blots were performed to validate the protein expression and confirm that reduced HR is not due to loss of protein expression. While RPP8 and BjWRR1 lacked bands for the MHD^MHV^ motif construct (figure 3, f, g) thus repeating the same pattern as Sr35 and RPM1 (figure 2, b), RPP13 overall lacked expression while still closely matching the HR-assay results of RPP8 (figure 3, f). This data shows that the preEDVID motif found in NLRs across the phylogeny of flowering plants could act in concert with the canonical EDVID motif to stabilise the CC-NLR packing. Thus, due to the nested position of the CC domain in the oligomer predictions that is reinforced by the preEDVID motif, NLRs with similarity to RPP8 may function in a similar manner to canonical NLRs like ZAR1 and Sr35.

**Figure 3.**
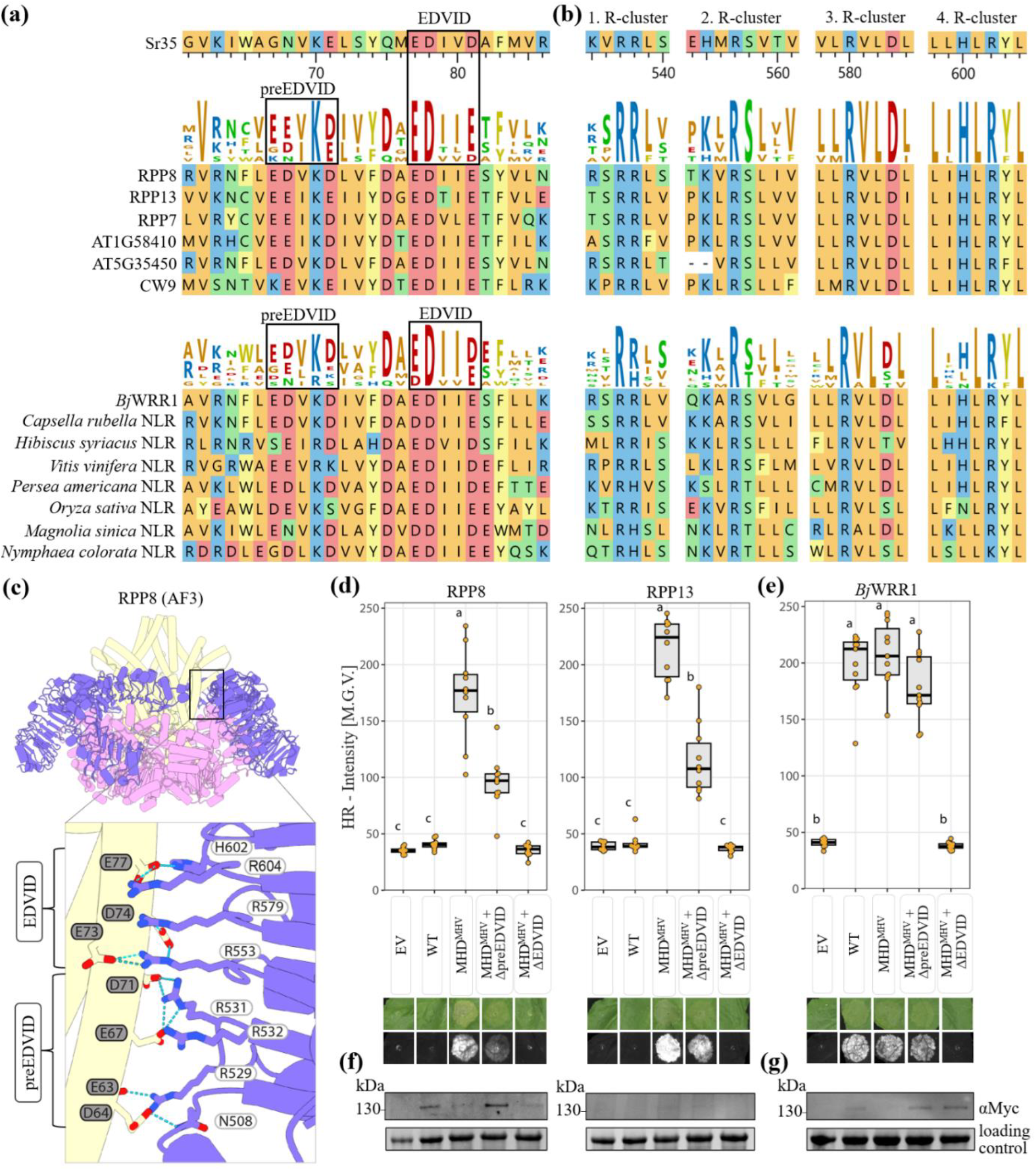
Acidic amino acids N-terminal of the canonical EDVID motif, called preEDVID motif, contribute to its function. **(a)** Alignments with Sr35 show high similarity in in Arabidopsis CC-NLRs a with a conserved EDVID motif and additionally conserved acidic residues N-terminally preceding it, called preEDVID motif. NCBI GenBank IDs are listed in supplementary table S2. **(b)** Alignments of the LRR blades show well conserved R-cluster. **(c)**, Salt bridges between the acidic residues of the EDVID and preEDVID motif and the R-cluster of the LRR-domain in RPP8 (AF3 prediction ipTM=0.71; pTM=0.73). **(d)** Hypersensitive response intensity of 5 weeks old *Nicotiana benthamiana* measured through infrared fluorescence at 800 nm with the iBright® imaging system. Displayed are the mean grey values of the cell death measured 4 days post infiltration with *Agrobacterium tumefaciens*. Substituted amino acids for RPP8, RPP13 and *Bj*WRR1 are listed in table 1. The box plots display the median and interquartile range; n=10; different letters represent significantly different groups of means, based on analysis of variance ANOVA and post-hoc Tuckey’s test P=<0.05. **(d)**, Western blot of α-Myc tagged NLR constructs expressed in *N. benthamiana*. Three replicates were pooled. Rubisco band on the blotting membrane as loading control.

To assess whether the EDVID motif can serve as a predictor of alternative CC domain topology in NLR oligomers, we examined the Arabidopsis NLRs which do not contain a clearly conserved EDVID sequence, also called the CC_G10_-type NLRs (Madhuprakash *et al*., 2024) Sequence alignment of Arabidopsis NLRs confirmed that the EDVID motif is less conserved in this group, exemplified by RPS2, which no longer contains any acidic residues at this site (figure 4, a). In parallel, we found that the region of the LRR domain typically enriched in basic residues in NLRs with a conserved EDVID motif contains fewer arginines, histidines, and lysines in SUMM2 and L5, particularly in the LRR blades 2 and 3 (figure 4, b). However, L5 and SUMM2 retain acidic residues in the region that corresponds to the EDVID motif in NLRs with the conserved sequence (figure 4, a). To investigate the functional significance of this region, we substituted the acidic residues in L5 and SUMM2 and tested their activity in Nicotiana HR assays. While WT SUMM2 did not induce HR beyond the vector control, WT L5 triggered a strong HR response, suggesting a lack of negative regulation in the heterologous system (figure 4, c). Both SUMM2 and L5 produced a robust HR phenotype when carrying the autoactivating MHV substitution (figure 4, c). Notably, replacement of the acidic residues in the EDVID-corresponding region had no detectable effect on HR activity, as the MHD^MHV^ ΔEDVID double motif mutants displayed responses similar to their respective MHD^MHV^ motif mutants (figure 4, c). A validation of the protein expression in Nicotiana was performed through Western blots (figure 4, d). While for SUMM2 even the MHD^MHV^ construct band is visible, the autoactive L5 only shows a faint band for the WT that is completely diminished for constructs with the MHD^MHV^ background that produce a high cell death (figure 4, d). We then analysed the structural context of these residues using the predicted oligomer models (figure 4, e). In both proteins, the relevant acidic residues, E83 and E88 in SUMM2 and D87 and D89 in L5, are located within the α3-helix of the CC domain, but this region is exposed to the solvent and does not appear to mediate stabilizing interactions with the LRR domain (figure 4, e). These findings suggest that the canonical EDVID motif, along with its role in mediating interactions with the LRR domain, has been lost in these CC_G10_-type NLRs. As a result, NLRs lacking the conserved EDVID motif may display a structurally divergent CC domain topology that functions independent of EDVID-R-cluster structural packing.

**Figure 4.**
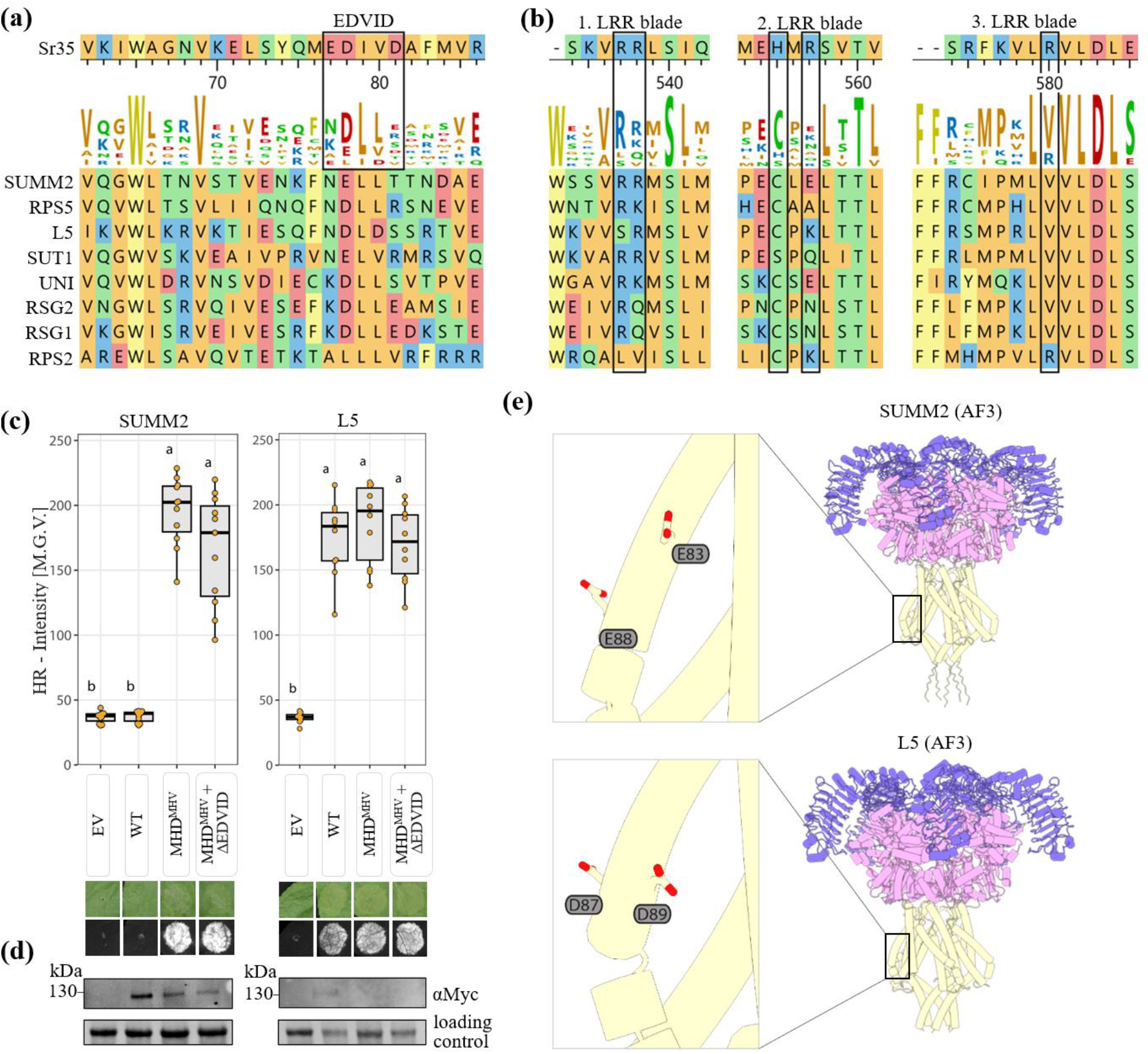
Acidic residues in the α3-helix of NLRs without conserved EDVID motif are dispensable for CC domain function. **(a)** Alignments with Sr35 show few polar amino acids in the corresponding EDVID area. **(b)** Alignments show less conserved R-cluster of CC_G10_-NLRs compared to Sr35. **(c)** Hypersensitive response intensity of 5 weeks old *Nicotiana benthamiana* measured through infrared fluorescence at 800 nm with the iBright® imaging system. Displayed are the mean grey values of the cell death measured 4 days post infiltration with *Agrobacterium tumefaciens*. The box plots display the median and interquartile range; n=10; different letters represent significantly different groups of means, based on analysis of variance ANOVA and post-hoc Tuckey’s test P=<0.05. **(d)**, Western blot of α-Myc tagged NLR constructs expressed in *N. benthamiana*. Three replicates were pooled. Rubisco band on the blotting membrane as loading control. **(e)** No contact between the LRR-and CC-domain was shown in the AF3 prediction of SUMM2 (ipTM=0.60; pTM=0.63) and L5 (ipTM=0.51; pTM=0.56).

Structure prediction of NRG1.1 using AlphaFold 3 revealed a CC domain topology resembling that of SUMM2 and L5, characterised by a CC barrel protruding from the NBD side. However, the confidence scores and expected position error for this conformation in NRG1.1 were notably low (figure 1, b, c). Sequence alignment of the α3-helix containing the EDVID motif in canonical NLRs such as Sr35 and ZAR1, along with the lateral LRR region harbouring the R-cluster, showed that while NRG1.1 retains weak conservation of the EDVID motif, the R-cluster is comparatively well conserved (figure 5, a, b). To assess the functional relevance of acidic residues in the EDVID-corresponding region of NRG1.1, we generated amino acid substitutions and tested their activity in Nicotiana hypersensitive response assays. Due to low sequence similarity between the CC domains of Sr35 and NRG1.1, it was difficult to precisely identify the EDVID-equivalent region based on sequence alone. Therefore, we used the x-ray crystal structures of two mutant NRG1.1 CC_R_ domains of (Jacob *et al*., 2021) as a structural guide. Two acidic clusters within this helix were identified and designated EDVID 1 (D63, E68, E71) and EDVID 2 (D74, E77, D80). Amino acid substitutions in these clusters were introduced into the autoactive NRG1.1 MHV background, which displays strong HR (figure 5, c). While the MHD^MHV^ ΔEDVID 2 double motif mutant retained robust HR activity, the MHD^MHV^ ΔEDVID 1 double motif mutant completely lost HR function, indicating that the residues in the EDVID 1 motif are essential for NRG1.1-mediated cell death. This result is supported by sequence alignment of NRG1.1 homologs across diverse plant lineages, which showed strong conservation of D63 and E68 within EDVID 1, in contrast to the weak conservation of EDVID 2 residues (figure 5, a). The Western blot of the NRG1.1 constructs validated their expression in Nicotiana (figure 5, d). Although AlphaFold 3 predicts a protruded CC domain topology for NRG1.1 (figure 5, e), this model is associated with particularly low confidence (figure 1, d). In this structural prediction, D63 and E68 are positioned on the surface of the CC barrel. However, given their essential role in HR activity and evolutionary conservation, this surface-exposed position may not reflect their functional conformation suggesting that the model does not accurately capture the physiologically relevant oligomeric structure of NRG1.1 in a cellular context. The function of the EDVID 1 motif in the CC_R_-NLRs remains to be demonstrated through a empirically determined structure of these proteins.

**Figure 5.**
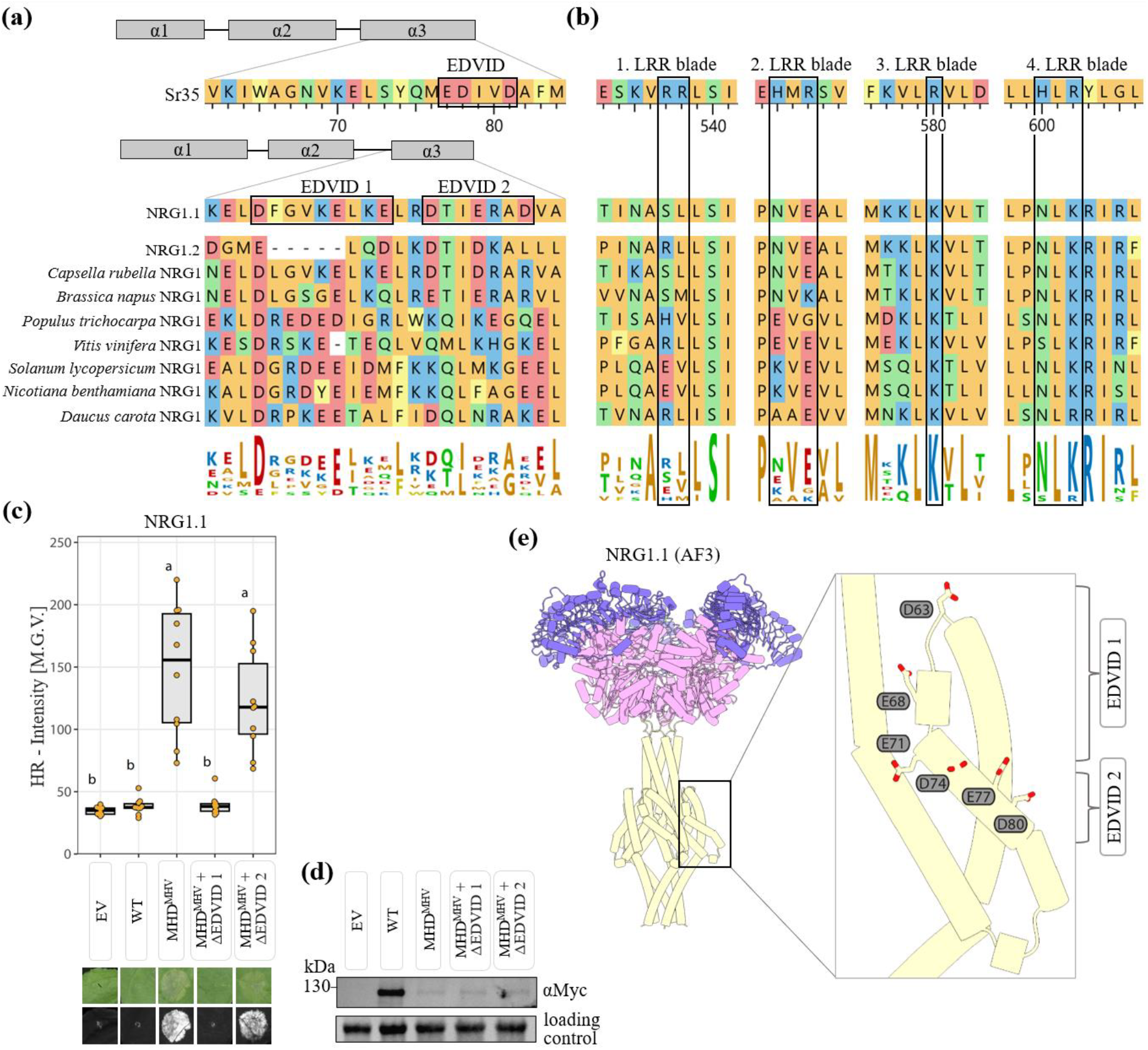
NRG1.1 shows a loss-of-function upon mutation of acidic residues in the EDVID-corresponding area in the α3-helix of the CC domain. **(a)** Alignments with Sr35 shows no conserved EDVID motif, but few conserved acidic residues in NRG1 and homologs of different species. GenBank IDs are displayed in supplementary table S2. **(b)** Alignments with Sr35 show partially conserved R-cluster beginning at the third LRR-blade. **(c)** Hypersensitive response intensity of 5 weeks old *Nicotiana benthamiana* measured through infrared fluorescence at 800 nm with the iBright® imaging system. Displayed are the mean grey values of the cell death measured 4 days post infiltration with *Agrobacterium tumefaciens*. The box plots display the median and interquartile range; n=10; different letters represent significantly different groups of means, based on analysis of variance ANOVA and post-hoc Tuckey’s test P=<0.05. **(c)** No contact between the LRR and CC domain was shown in the AF3 prediction of NRG1.1 (ipTM=0.57; pTM=0.60). **(d)** Western blot of α-Myc tagged NLR constructs expressed in *N. benthamiana*. Three replicates were pooled. Rubisco band on the blotting membrane as loading control.

## Discussion

Our study provides new insights into the regulatory roles of the EDVID motif in NLR cell death activity. Across a diverse panel of Arabidopsis CC-NLRs, we find that conservation of the EDVID motif robustly predicts a canonical CC-LRR topology resembling that of ZAR1 and Sr35. This includes the nested positioning of the CC domain barrel, anchored via salt bridges between acidic EDVID residues and basic residues on the LRR domain. These findings support the hypothesis that the EDVID motif functions as a structural determinant of canonical resistosome architecture and suggest that it plays a stabilizing role in securing domain orientation necessary for NLR activity.

### Role of EDVID in CC domain regulation

Due to its unique three-dimensional position in certain CC-NLRs, the EDVID motif serves as a critical physical contact between the signalling CC domain and the pathogen-recognizing LRR domain. In the inactive state of ZAR1, it forms the only direct connection between these two domains, whereas in resistosomes such as ZAR1, Sr35, *Nb*NRC2 and *Sl*NRC4 ((Wang *et al*., 2019a; Förderer *et al*., 2022; Madhuprakash *et al*., 2024; Liu *et al*., 2024)), the packing function of the EDVID motif is supported via several structural redundancies. These structural redundancies include a tight packing of protomers in the oligomer and structural contacts between the NOD module and the CC domain, which collectively stabilise the CC barrel for cell death initiation. We hypothesise, that the EDVID motif has a dual function of providing regulatory control in the inactive receptor and in contributing to structural stability of the CC domain in the active resistosome.

Evidence for a regulatory role in the inactive state comes from several observations. For instance, deletion of the LRR domain in an MHD-activated form of Pm21 fails to trigger cell death (Gao *et al*., 2020) Similarly, in the NLR Rx, mutation of the EDVID motif disrupts binding and impairs CC-mediated cell death activity in response to activation by the PVX coat protein via the LRR domain. Our own data further support this model, showing that EDVID mutations in NLRs such as RPP8, RPP13, RPM1, and *Bj*WRR1 abolish cell death activity in MHD-activated backgrounds. Similarly, the EDVID motif is also required in *Sl*NRC4 for the function of an MHD-activated mutant, as shown by a loss of activity in an MHD^MHV^ ΔEDVID double motif mutant of *Sl*NRC4 (Liu *et al*., 2024). Together, these findings suggest that the EDVID motif acts as a ‘mechanical hinge’, enabling the transfer of torque from the pathogen detecting LRR domain, through the NOD module, to the CC domain that mediates downstream signalling. This highlights the importance of the CC-LRR contact in relaying the conformational state of the NOD module to the CC domain.

In contrast, the activity retention of our Sr35 EDVID mutant in the context of autoactivation by MHD substitution prompts questions regarding MHD-mutation versus effector-mediated activation. This could be explained by additional interactions such as between NOD and CC domains that may compensate for the loss of the CC-LRR contact and that collectively stabilise the CC barrel for cell death initiation in Sr35. However, it remains unclear why these additional structural features fail to retain CC domain activity in MHD^MHV^ ΔEDVID mutants of other NLRs such as RPM1 (our data) and *Sl*NRC4 (Liu *et al*., 2024).

In addition, slight shifts in the acidic residues of the EDVID and the basic residues of the R-cluster were observed from the comparison of the ZAR1 inactive versus the ZAR1 active form (Wang *et al*., 2019a,b) which underscores the fine-tuned regulation mediated by the EDVID motif. Careful dissection of the endogenous NLR activation mechanism including structure determination of resting and intermediate forms will be necessary to fully comprehend the evolutionary diversified NLR regulation. The diverged sensitivity of Sr35 to EDVID mutation in the MHD background may thus underscore particular receptor-specific distinctions. On the other hand, this finding also underpins the notion that MHD-mutation may trigger other activation trajectories in NLRs in comparison to activation by pathogens.

In this regard, our findings also contribute to a broader understanding of NLR activation dynamics. While the EDVID interface has been proposed to be temporarily disengaged during the CC fold-switching that occurs upon activation (Förderer *et al*., 2022), it may also act as a pre-activation stabiliser or molecular anchor, ensuring the correct orientation of domains for efficient signalling. Such anchoring could be critical for the mechanical coupling between pathogen recognition at the LRR and the conformational rearrangements in the NOD and CC domains required for resistosome formation and downstream signalling.

### An acidic preEDVID motif for enhanced signalling robustness

On the other hand, our finding that RPP8, RPP13 and *Bj*WRR1 MHD mutant are sensitive to EDVID mutation highlight the function of the EDVID for positive regulatory control of the CC domain exerted via CC-LRR contacts full-length receptors.

Next to this, mutations in a newly identified, upstream preEDVID motif caused a partial reduction in cell death. These results together with an extension of basic residues co-occurring in the LRR domain suggest an additive or cooperative function between EDVID and preEDVID motifs, supporting a layered mechanism of CC-LRR domain stabilisation. The presence of both motifs in NLRs with homology to RPP8, and their evolutionary distribution across derived and early angiosperm lineages, hints at selective pressure to maintain this structural reinforcement.

### Loss of EDVID motif in alternative NLR configurations

A structurally divergent mode of CC domain organisation was revealed in EDVID-lacking NLRs including SUMM2 and L5. In these proteins, AlphaFold 3 predicts a strikingly protruded CC domain conformation extending from the NOD side of the predicted oligomer, with no evident contact between the CC and LRR domains. While previous studies have proposed that members of this subgroup adopt distinct oligomerisation modes, referred to collectively as CC_G10_-type NLRs (Madhuprakash *et al*., 2024), our data provide functional support for this distinction: EDVID-associated activity, critical in canonical NLRs, is dispensable in these receptors. Next to SUMM2 and L5, the receptor RPS2 belongs to the same group and has lost the EDVID motif from its sequence (Figure 4, a). Interestingly, the resting state of RPS2 was demonstrated to have biochemical properties of an integral membrane protein (Axtell & Staskawicz, 2003). Together, these data support a divergent architecture of EDVID-lacking NLRs in both inactive and potentially active states, suggesting that they represent a mechanistically distinct subclass of these immune receptors.

Despite the growing utility of structure prediction tools included in this study, important limitations remain. For NRG1.1, AlphaFold 3 models predict a protruded CC domain topology with notably low confidence scores, casting doubt on their physiological relevance. Supporting this uncertainty, recent cryo-EM structures of the helper NLRs NRG1A and NRG1C in complex with EDS1-SAG101 (Huang *et al*., 2025) resolved the LRR-WHD region but did not capture the CC domain, leaving open the question of whether a CC-LRR interface is formed in these complexes. While functional evidence underscores that the NRG1 CC domain alone is sufficient to trigger cell death (Collier *et al*., 2011), our own assays further identified a conserved acidic cluster within the α3-helix of NRG1.1 as essential for cell death in the context of the full-length, MHD-activated NLR. These findings suggest that the topology of the CC domain in an NRG1.1 oligomeric context depends on structural features not captured in current in silico models. High-resolution structural studies of full-length NRG1 oligomers, particularly in membrane-associated states, will be essential to resolve this discrepancy and elucidate the mechanism of NRG1-mediated immunity and activation.

Taken together, our results position the EDVID motif as both a structural hallmark and functional predictor of canonical NLR architecture. Its presence delineates a mechanistic subclass of NLRs, that is typified by ZAR1, Sr35, RPM1, and RPP8 and that likely share conserved modes of oligomerisation and activation. Conversely, the absence of the EDVID motif found in CC_G10_ and helper NLRs correlates with distinct structural arrangements and possibly non-canonical modes of signalling.

