## Supplementary Tables for "Diversification of the “EDVID” packing motif underpins structural and functional variation in plant NLR coiled-coil domains"

### Supplementary Materials

Supplementary Table S1: List of all Arabidopsis CC-NLR originally clustered by (Wróblewski et al., 2018). The NLRs relevant to this study highlighted.

| Phylogenetic Groups | Groups of Wróblewski et al., 2018 | Representative Gene Model Name | Primary Gene Symbol |
|---|---|---|---|
| $CC_R$ | A | AT1G33560.1 | ACTIVATED DISEASE RESISTANCE 1 (ADR1) |
| $CC_R$ | A | AT4G33300.1 | ADR1-LIKE 1 (ADR1-L1) |
| $CC_R$ | A | AT5G04720.1 | ADR1-LIKE 2 (ADR1-L2) |
| $CC_R$ | A | AT5G66630.1 | DA1-RELATED PROTEIN 5 (DAR5) |
| $CC_R$ | **A** | **AT5G66900.1** | **N REQUIREMENT GENE 1.1 (NRG1.1)** |
| $CC_R$ | A | AT5G66910.1 | N REQUIREMENT GENE 1.2 (NRG1.2) |
| $CC_{G10}$ | B | AT1G12210.1 | RPS5-LIKE 1 (RFL1) |
| $CC_{G10}$ | B | AT1G12220.1 | RESISTANT TO P. SYRINGAE 5 (RPS5) |
| $CC_{G10}$ | **B** | **AT1G12280.1** | **SUPPRESSOR OF MKK1 MKK2 2 (SUMM2)** |
| $CC_{G10}$ | **B** | **AT1G12290.1** | **(L5)** |
| $CC_{G10}$ | B | AT1G15890.1 | |
| $CC_{G10}$ | B | AT1G51480.1 | RESISTANCE SILENCED GENE 1 (RSG1) |
| $CC_{G10}$ | B | AT1G61180.2 | (UNI) |
| $CC_{G10}$ | B | AT1G61190.1 | |
| $CC_{G10}$ | B | AT1G61310.1 | |
| $CC_{G10}$ | B | AT1G62630.1 | |
| $CC_{G10}$ | B | AT1G63350.2 | |
| $CC_{G10}$ | B | AT1G63360.1 | |
| $CC_{G10}$ | B | AT4G10780.1 | |
| $CC_{G10}$ | B | AT4G14610.1 | |
| $CC_{G10}$ | B | AT4G26090.1 | RESISTANT TO P. SYRINGAE 2 (RPS2) |
| $CC_{G10}$ | B | AT4G27190.1 | |
| $CC_{G10}$ | B | AT5G05400.1 | |
| $CC_{G10}$ | B | AT5G43730.1 | RESISTANCE SILENCED GENE 2 (RSG2) |
| $CC_{G10}$ | B | AT5G43740.1 | |
| $CC_{G10}$ | B | AT5G47250.1 | |
| $CC_{G10}$ | B | AT5G47260.1 | |
| $CC_{G10}$ | B | AT5G63020.1 | SUPPRESSORS OF TOPP4-1 (SUT1) |
| canonical EDVID | C | AT1G50180.1 | CEL-ACTIVATED RESISTANCE 1 (CAR1) |
| canonical EDVID | **C** | **AT3G07040.1** | **RESISTANCE TO P. SYRINGAE PV MACULICOLA 1 (RPM1)** |
| canonical EDVID | C | AT3G14460.1 | LEUCINE-RICH REPEAT (LRR) PROTEIN 1 (LRRAC1) |
| canonical EDVID | C | AT3G14470.1 | LEUCINE RICH REPEAT PROTEIN 1 (LRR4) |

| | | | |
|---|---|---|---|
| canonical EDVID | C | AT3G46530.1 | RECOGNITION OF PERONOSPORA PARASITICA 13 (RPP13) |
| canonical EDVID | C | AT3G46710.1 | |
| canonical EDVID | C | AT3G46730.1 | |
| canonical EDVID | C | AT3G50950.1 | HOPZ-ACTIVATED RESISTANCE 1 (ZAR1) |
| canonical EDVID + preEDVID | D | AT1G53350.1 | |
| canonical EDVID + preEDVID | D | AT1G58390.1 | |
| canonical EDVID + preEDVID | D | AT1G58400.2 | |
| canonical EDVID + preEDVID | D | AT1G58410.1 | |
| canonical EDVID + preEDVID | D | AT1G58602.1 | RECOGNITION OF PERONOSPORA PARASITICA 7 (RPP7) |
| canonical EDVID + preEDVID | D | AT1G58807.1 | |
| canonical EDVID + preEDVID | D | AT1G58848.1 | |
| canonical EDVID + preEDVID | D | AT1G59124.2 | |
| canonical EDVID + preEDVID | **D** | **AT1G59218.1** | **RPP13** |
| canonical EDVID + preEDVID | D | AT1G59620.1 | (CW9) |
| canonical EDVID + preEDVID | D | AT1G59780.1 | |
| canonical EDVID + preEDVID | D | AT5G35450.1 | |
| canonical EDVID + preEDVID | **D** | **AT5G43470.1** | **RECOGNITION OF PERONOSPORA PARASITICA 8 (RPP8)** |
| canonical EDVID + preEDVID | D | AT5G48620.1 | |
| excluded | E | AT1G10920.1 | LOCUS ORCHESTRATING VICTORIN EFFECTS1 (LOV1) |
| excluded | E | AT1G52660.1 | |
| excluded | E | AT1G61300.1 | |
| excluded | E | AT3G15700.1 | |
| excluded | E | AT4G19060.1 | |
| excluded | E | AT4G27220.2 | |
| excluded | E | AT5G45440.1 | |
| excluded | E | AT5G45490.1 | |

Supplementary Table S2: NRG1.1 and RPP8-preEDVID homologs that were used in alignments.

| Phylogenetic Groups | NCBI GenBank ID | Organism | Primary Gene Symbol |
|---|---|---|---|
| **canonical EDVID + preEDVID** | **MK469977.1** | ***Brassica juncea*** | **WRR1** |
| canonical EDVID + preEDVID | XP_006303146.1 | *Capsella rubella* | probable disease resistance RPP8-like protein 2 isoform X1 |
| canonical EDVID + preEDVID | VEPZ02000817.1 | *Hibiscus syriacus* | RPP8 |
| canonical EDVID + preEDVID | DRVX08091.1 | *Vitis_vinifera* | putative disease resistance RPP8-like protein 4 |
| canonical EDVID + preEDVID | KAJ8646342.1 | *Persea americana* | hypothetical protein MRB53_008090 |
| canonical EDVID + preEDVID | EAY96750.1 | *Oryza sativa (Indica Group)* | hypothetical protein OsI_18670 |
| canonical EDVID + preEDVID | XP_058107017.1 | *Magnolia sinica* | putative disease resistance protein_RGA3 |
| canonical EDVID + preEDVID | XP_031482955.1 | *Nymphaea colorata* | putative disease resistance protein_RGA3 |
| $CC_R$ | XP_006281815.1 | *Capsella rubella* | probable disease resistance protein At5g66900 |
| $CC_R$ | XP_022568565.1 | *Brassica napus* | probable disease resistance protein At5g66900 |
| $CC_R$ | XP_024461167.2 | *Populus trichocarpa* | probable disease resistance protein At5g66900 |
| $CC_R$ | XP_002268207.3 | *Vitis vinifera* | probable disease resistance protein At5g66900 |
| $CC_R$ | XP_004232056.1 | *Solanum lycopersicum* | probable disease resistance protein At5g66900 |
| $CC_R$ | AAY54606.1 | *Nicotiana benthamiana* | NRG1 |
| $CC_R$ | XP_017243250.1 | *Daucus carota* | probable disease resistance protein At5g66900 |
